## Supplemental Text for Pech et al for "Synaptic deregulation of cholinergic projection neurons causes olfactory dysfunction across 5 fly Parkinsonism models"

**Supplemental data for Pech et al.**

The supplemental data consists of 8 Supplemental files and 4 Supplemental figures.

**Supplemental file 1:** List of gBlocks and oligos used to generate and validate the flies generated in this study. List of full fly genotypes used in this study.

**Supplemental file 2:** Full results of the Cell Ranger summaries (10xGenomics).

**Supplemental file 3**: Summary of the number of cells per cell cluster for each mutant and controls, including the raw number of cells, the percentage of cells for each cluster per mutant and the percentage of cells for each mutant and controls per cluster.

**Supplemental file 4**: Summary of the parameters used to model DEG-cell number correlation.

**Supplemental file 5**: List of cell types and respective z-residuals (deviation from model) to determine affected cell types and those that are outside of the 95% confidence interval.

**Supplemental file 6:** List of the clusters with significant changes (based on cell number-adjusted DEGs) for each model, black cell clusters (cells with significant number of DEG) are listed as TRUE.

**Supplemental file 7**: Summarized results of the DEG analysis in fly model brains and post-mortem human samples and the summary of the SynGO analysis.

**Supplemental file 8:** List of software versions used in this study.

**Supplemental Figure legends**

**Supplemental Fig. 1. Characterization of PD knock-in models,** *related to Figure 1.*

(**a**) RNA expression levels, measured by RT-PCR with primers to the indicated CDS (above the graphs) in heads of controls and animals with wild-type or pathogenic mutant CDS knocked-in. Data are expressed as percent of endogenous Drosophila gene expression. Grey bars indicate the variance in controls. (**b**) Survival curves of flies kept under wet starvation (See median survival in Fig. 1). (**c**) Example images of mitochondrial TMRE fluorescence at larval NMJs.

Insets: magnified synaptic boutons (See also Fig. 1). Scale bar: 10 µm. (**d**) Example electroretinogram (ERG) recordings (1 s light stimulus); See also Fig. 1. Animals were kept for 7 days in constant light prior to the experiment. Red line: the average of the individual raw data traces that are shown.

**Supplemental Fig. 2. Validation and analysis of single-cell RNA sequencing data,** *related to Figure 2***.**

(**a**) Frequency of individual cell types (olfactory projection neurons (OPN), dopaminergic neurons (DAN), visual projection neurons (T1)) and the sex ratios of the sequenced cells detected across the PD knock-in models and controls. Note, we only sequenced males from *hPINK1*$^{P399L}$ mutants. Error bars represent mean ± 95 % CI. (**b**) Alignment of published transcriptomic profiles defined by bulk sequencing of FAC sorted cell types (y-axis, see also (Davie et al., 2018)) and the cell types identified in this study (x-axis), validating the identification of cell types in the tSNE clusters. Green indicates central brain cell types; Blue is optic lobe; Yellow: glia; Grey: others. (**c**) Expression level of knocked-in CDS across the cell types, indicating the knocked-in genes are broadly expressed. DAN - dopaminergic neurons (PAM cluster), OPN - olfactory projection neurons, T1 - T1 visual interneurons. (**d**) Comparison of two different algorithms to detect cell-type specific DEG (See methods) showing a similar number of DEG and high correlation of signed p-values. (**e**) Graphs showing the number of DEG versus the number of cells in a cluster (each dot is a cell type) for each PD fly model and the model of cell-number-dependency of p-values (dotted line, See methods). Confidence intervals in red. Cell types with residuals above confidence intervals are considered affected.

**Supplemental Fig. 3. Single-cell RNA sequencing of cholinergic regions in post-mortem human brain samples,** *related to Figure 3***.**

(**a**) tSNE of cells characterized in three cholinergic brain regions of ten brains, annotated are the major cell types (left) and 36 individually identified cell types (numbered, right). N - neurons, OG - oligodendrocytes, ENT - endothelial cells, EPY - ependymal cells, LM - lymphocyte, MG - microglia, AST - astrocytes, OPC - oligodendrocyte precursor cells, U - unknown. (**b**) Frequency of the 36 cell types in pooled PD vs. non-PD samples, in individual brain samples, and in the three brain regions. (**c**) tSNE, as shown in (a), with different neuronal marker gene expression indicated (counts of Unique Molecular Identifier, UMI). (**d**) Frequency of cells expressing cholinergic marker genes *ACHE* and *CHAT*. NBM - *Nucleus basalis Meynert*, Nac - *Nucleus accumbens*, Put - *Putamen*.

**Supplemental Fig. 4. Generation and validation of human LRRK2$^{G2019S}$ ventral midbrain DAN,** *related to Figure 6***.** (**a**) Schematic representation of gene editing strategy to knock-in the G2019S mutation in the *LRRK2* locus. (**b**) Sanger sequencing of a single gene edited clone showing successful homozygous editing of the indicated nucleotide. (**c**) KOLF2-1J wildtype (LRRK2$^{WT}$) control iPSCs and KOLF2-1J LRRK2$^{G2019S/G2019S}$ (LRRK2$^{G2019S}$) iPSCs show normal expression of pluripotency markers OCT4 (yellow), SOX2 (purple), NANOG (purple) and TRA-1-81 (green). Nuclei are counterstained with DAPI (blue). Scale bar: 100 µm. (**d**) KOLF2-1J wildtype (LRRK2$^{WT}$) control and KOLF2-1J LRRK2$^{G2019S/G2019S}$ (LRRK2$^{G2019S}$) mutant ventral midbrain neural progenitor cells show normal expression of FOXA2 (magenta), LMX1A (green) and OTX2 (green), confirming that the neural progenitor cells are ventral midbrain-specific and capable of differentiating into DAN. Nuclei are counterstained with DAPI (blue). Scale bar: 100 µm. (**e**) Quantification of the frequency of TH+/MAP2+ DAN. Statistical significance calculated with an ordinary T-test: ns, not significant. Error bars represent mean ± SEM. Data were collected in three independent vmDAN differentiations. (**f**) Differentiation protocol used.
