## Supplementary figures and images for "Synaptic deregulation of cholinergic projection neurons causes olfactory dysfunction across 5 fly Parkinsonism models"

### Supplemental Figure 1

#### Supplemental Figure 1

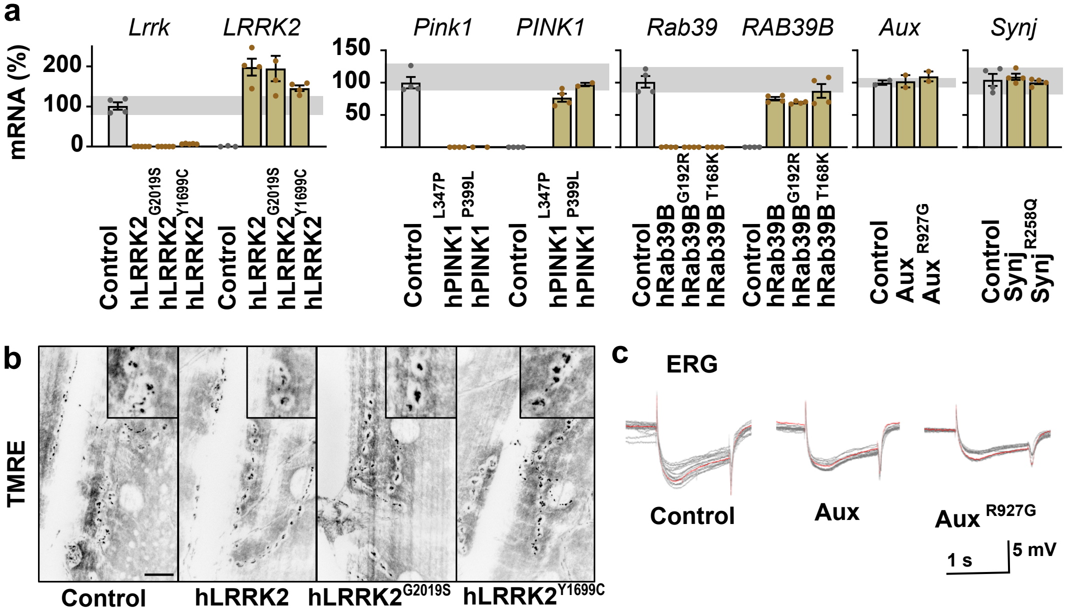

### Supplemental Figure 2

Supplemental Figure 2

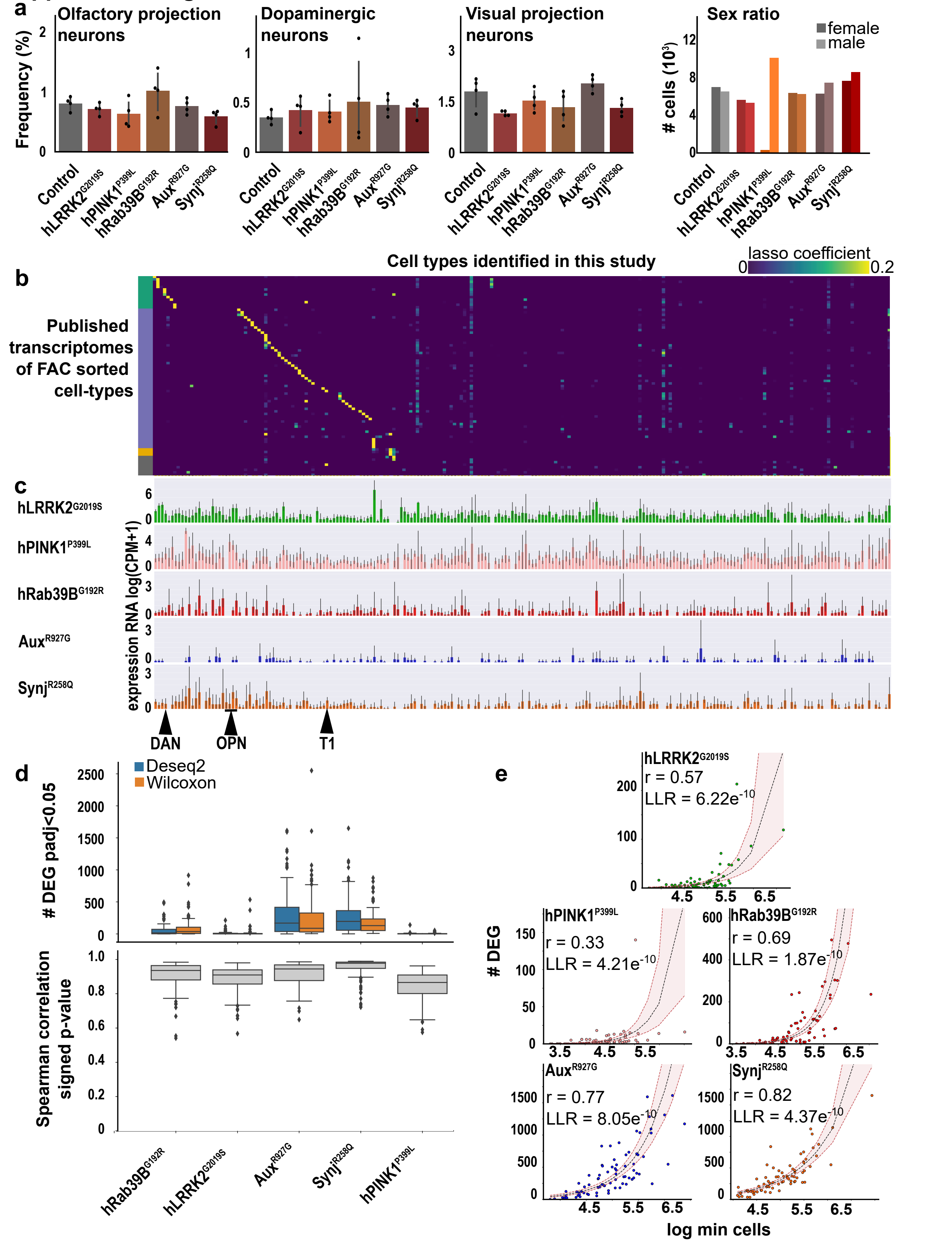

### Supplemental Figure 3

##### Supplemental Figure 3

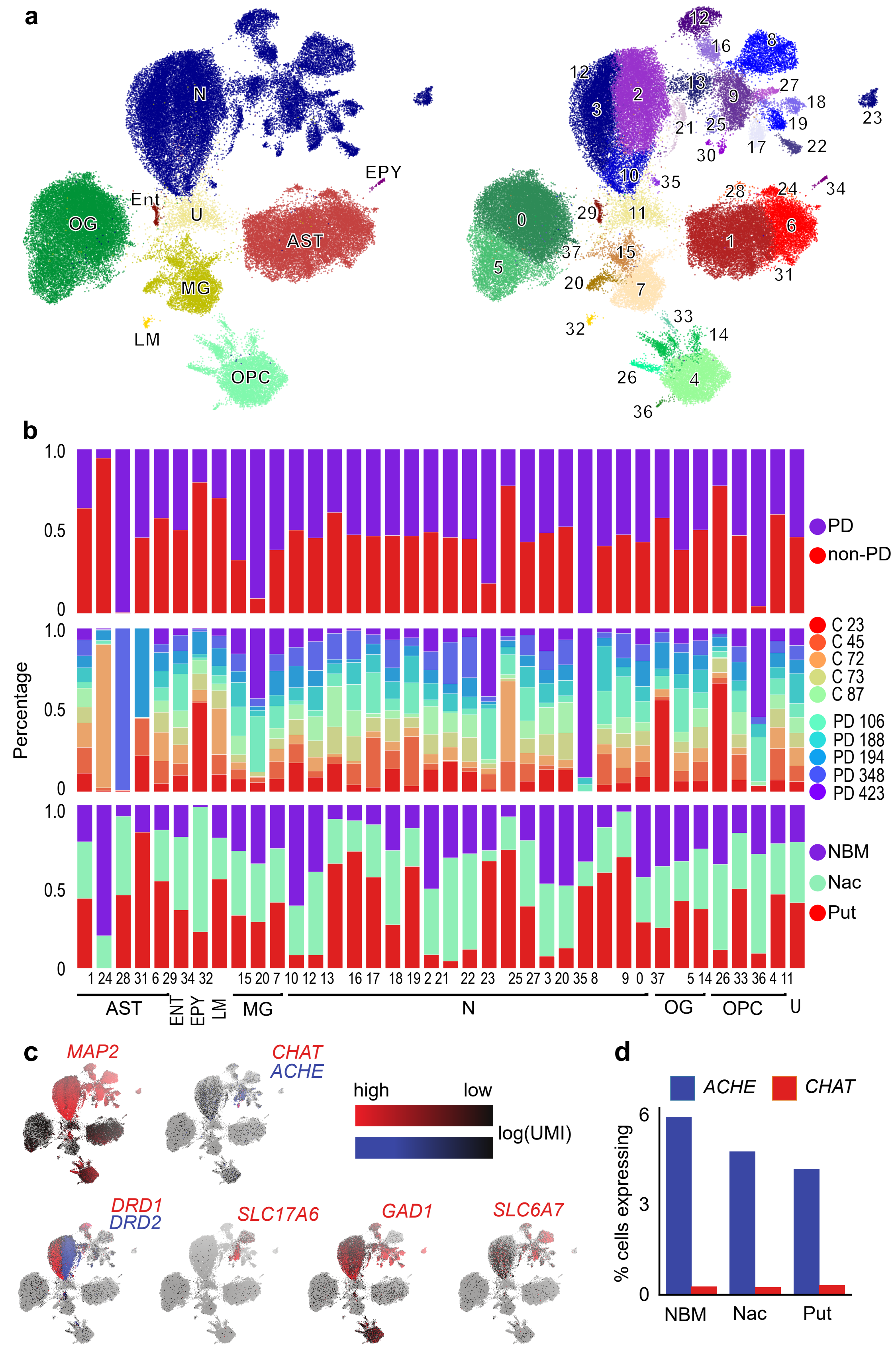

### Supplemental Figure 4

#### Supplemental Figure 4

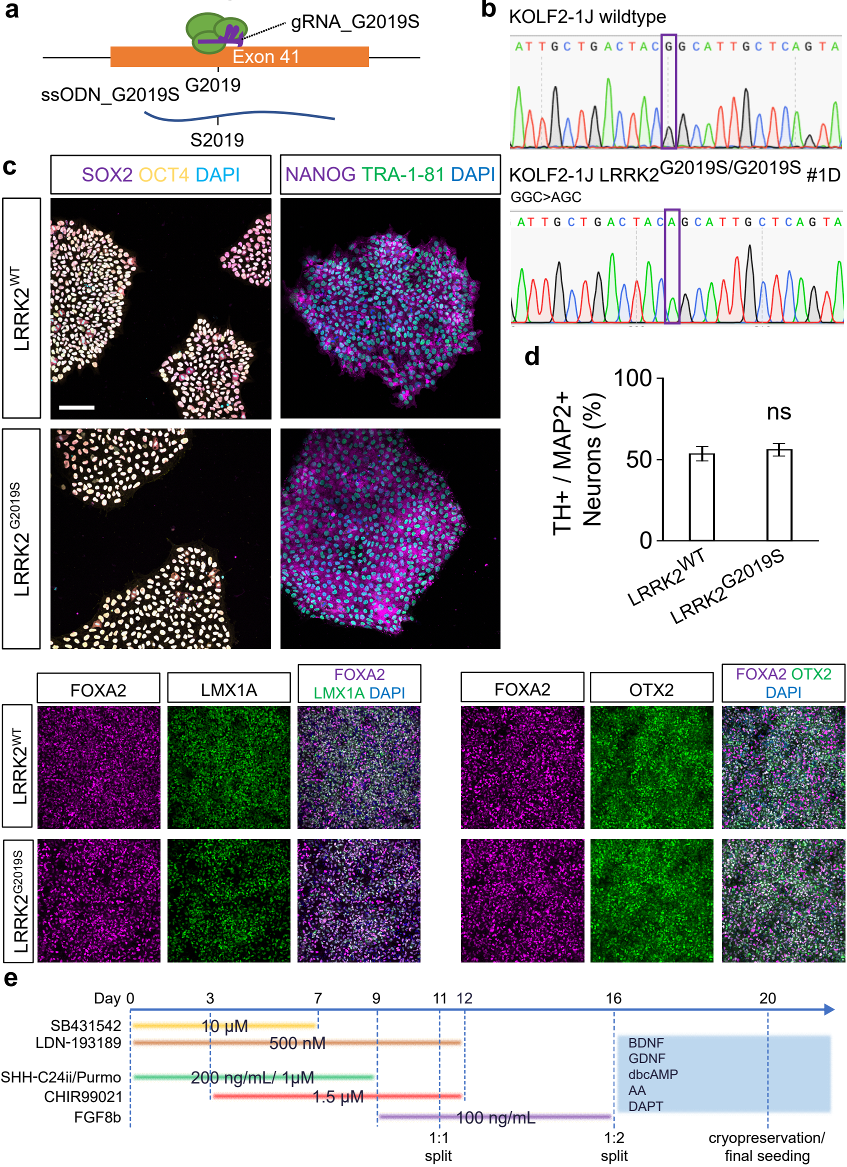
